# GeneReL: A Large Language Model-Powered Platform for Gene Regulatory Relationship Extraction with Community Curation

**DOI:** 10.64898/2026.02.10.705020

**Authors:** Joo-Seok Park, Su-Yeon Ha, Yejin Lee, Yang Jae Kang

## Abstract

**Motivation:** Gene regulatory networks provide fundamental insights into plant biology, yet extracting structured interaction data from scientific literature remains a significant bottleneck. Traditional manual curation cannot scale to meet the demands of modern research, while automated text mining approaches struggle with the complexity of gene nomenclature and relationship classification. Large language models offer promising capabilities for information extraction, but integrated platforms combining LLM extraction with community validation for plant regulatory databases remain scarce.

**Results:** We developed GeneReL, an integrated platform combining LLM-based extraction with community-driven curation for gene regulatory networks in *Arabidopsis thaliana*. The system employs a tiered pipeline using Claude Haiku 4.5 for screening, Claude Sonnet 4 for extraction, and Claude Opus 4 for verification, along with a novel five-step gene normalization pipeline incorporating paper-text search and LLM-based disambiguation with UniProt annotations. The database contains 13,710 curated interactions across 51 relationship types, with 90.2% classified as high confidence based on linguistic certainty markers in source text. Comparison with IntAct reveals 86.8% of interactions are unique to our literature-derived database, demonstrating complementary coverage to existing resources. The web platform provides card-based browsing with voting capabilities, interactive network visualization using Cytoscape.js with locus-ID-based node consolidation, and administrative interfaces for curator review of ambiguous gene mappings.

**Availability and Implementation:** GeneReL is freely accessible at https://generel.newgenes.me.

## 1 Introduction

Gene regulatory networks (GRNs) represent the complex web of molecular interactions governing cellular processes in plants. In *Arabidopsis thaliana*, decades of research have elucidated numerous regulatory pathways controlling flowering time, organ identity, hormone signaling, and stress responses [1]. The importance of comprehensive GRN databases has driven development of resources such as AGRIS and specialized tools like AGENT for exploring published regulatory networks [2,3]. However, significant challenges remain in capturing the rapidly growing body of regulatory information scattered across thousands of publications.

### 1.1 The Information Extraction Challenge

The biomedical literature expands at an unprecedented rate, with gene-gene interactions typically described in natural language text. Traditional manual curation approaches cannot scale proportionally with literature growth—a systematic analysis of TAIR curation workflows estimated that only approximately 30% of relevant *Arabidopsis* literature receives curator attention [4]. Community curation has been explored as a strategy, with the CACAO project demonstrating that undergraduates can contribute high-quality functional annotations [5]. However, even successful community initiatives require substantial coordination and expertise.

Large language models (LLMs) represent a paradigm shift in text processing capabilities, with growing applications in plant biology [6]. Recent evaluations have benchmarked LLMs including GPT-4 and domain-specific models on biomedical tasks such as named entity recognition and relation extraction [7,8]. A comprehensive evaluation of 21 LLMs on gene regulatory relation extraction found that GPT-4 and Claude-Pro achieved F1 scores of 0.4448 and 0.4386 respectively, demonstrating both potential and remaining challenges for this task [9]. These studies reveal that LLMs can achieve competitive performance compared to traditional methods, particularly for complex information extraction requiring contextual understanding [10]. Recent work has also explored LLMs for gene regulatory network inference from expression data (LLM4GRN), though literature-based extraction remains underexplored [11]. The GIX framework demonstrated a three-stage approach for automated extraction of genetic interactions from literature using language models [12], while TRRUST pioneered sentence-based text mining for TF-target interaction curation in human and mouse [13].

### 1.2 Distinction from Existing Resources

Several databases provide regulatory interaction data for plants. AGRIS (Arabidopsis Gene Regulatory Information Server) and its successor AtRegNet represent the primary *Arabidopsis*-specific regulatory network resources, containing transcription factor binding site information and curated TF-target interactions [14]. PlantRegMap extends regulatory mapping across 165 plant species with curated TF-target interactions derived from ChIP-seq and other high-throughput experiments [15]. PlantTFDB provides comprehensive transcription factor annotations with regulatory relationship predictions [16].

GeneReL complements these resources by: (1) extracting relationships beyond TF-target interactions to include post-translational modifications, physical interactions, and miRNA targeting (51 distinct relationship types vs. 3-4 in AGRIS); (2) capturing regulatory evidence from narrative text rather than high-throughput data alone; and (3) implementing community validation through voting mechanisms for continuous quality improvement.

RegNetwork 2025 recently demonstrated large-scale literature-based regulatory network extraction for mammalian systems, integrating 31 databases and over 20,000 published papers to compile more than 11 million regulatory relationships for human and mouse [17]. GeneReL addresses the distinct needs of plant biology with *Arabidopsis*-specific gene nomenclature handling and a focus on semantic relationship types rather than association-based scoring.

Among broader molecular interaction databases, STRING integrates known and predicted protein-protein interactions across thousands of organisms. The 2025 update (STRING v12.0) now includes a regulatory interaction mode with directionality information, marking a significant enhancement [18]. IntAct provides expertly curated molecular interaction data derived from literature curation and high-throughput experiments, serving as a primary reference for protein interaction validation [19]. For *Arabidopsis*, STRING v12 contains 27,462 proteins with over 12.5 million putative interactions, though only 5.3% meet the high-confidence threshold (score >= 700).

### 1.3 Objectives

This study presents a platform integrating LLM-based extraction with community-driven curation for gene-gene interaction discovery. We address three objectives: (1) develop an automated pipeline leveraging the Claude API to extract structured regulatory relationships with gene identifier normalization; (2) implement a web-based platform for browsing, filtering, and visualizing interactions; and (3) establish community curation through voting mechanisms.

## 2 Materials and Methods

### 2.1 System Architecture

The platform comprises four integrated components: (1) data source management, (2) an automated extraction pipeline, (3) a Firebase backend for data storage and user management, and (4) a web frontend for visualization and community curation (**Supplementary Figure S1**).

### 2.2 Literature Retrieval

Full-text articles were retrieved from PubMed Central using the BioPython Entrez module. Publication date filters restricted retrieval to 2010-2024. PMC articles in JATS XML format were parsed using lxml (https://lxml.de/), with section-aware chunking employing overlapping segments of 3,000 words with 200-word overlap for articles exceeding the LLM context window.

### 2.3 LLM-Based Extraction

The extraction pipeline employs a tiered multi-model approach optimizing for both accuracy and cost-efficiency. We selected Claude models based on preliminary evaluations demonstrating stable extraction performance across varied scientific text scenarios, consistent with recent findings on LLM stability in chemical-disease extraction tasks [20]. Abstract screening uses Claude Haiku 4.5 (claude-haiku-4-5-20251001) to filter irrelevant papers before full extraction. Primary extraction uses Claude Sonnet 4 (claude-sonnet-4-20250514) based on its 200K token context window and strong performance on scientific text comprehension benchmarks. Verification uses Claude Opus 4 (claude-opus-4-20250514) to validate extracted interactions, confirming that both entities are genes (not phenotypes or processes), the evidence sentence supports the stated relationship, and the relationship type is correctly classified. The extraction prompt comprises task description, output schema definition, relationship taxonomy, and few-shot examples. Malformed JSON responses (approximately 2.3% of extractions) were re-queried with error feedback; persistent failures were logged and excluded (**Supplementary Table S1 and Supplementary Figure S2**).

Relationship types are constrained to a controlled vocabulary of 51 terms organized into eight categories: transcriptional regulation (e.g., Activates, Represses, Regulates, Modulates), physical interactions (e.g., Binds, Interacts, Recruits), post-translational modifications (e.g., Phosphorylates, Ubiquitinates, Methylates, Acetylates, SUMOylates), protein stability and processing (e.g., Stabilizes, Protects), post-transcriptional regulation (e.g., Targets), transport and localization (e.g., Transports, Translocates), genetic interactions (e.g., IsEpistaticTo, Antagonizes), and other relationships (e.g., Encodes, Synthesizes) (**Supplementary Table S2**).

Confidence levels are assigned based on linguistic markers: High for definitive experimental demonstration, Medium for indirect evidence, and Low for speculation or predictions. Evidence types classify experimental methodology (Genetic, Biochemical, Transcriptomic, ChIP, Y2H, BiFC, CoIP, Computational).

### 2.4 Gene Identifier Normalization

Gene names in scientific literature exhibit substantial heterogeneity, and gene symbols are not unique identifiers—one symbol may refer to multiple genes [21,22]. We developed a five-step hierarchical normalization pipeline to convert gene mentions to AGI locus format (**Figure 1; Supplementary Figure S3**):

**Figure 1.**
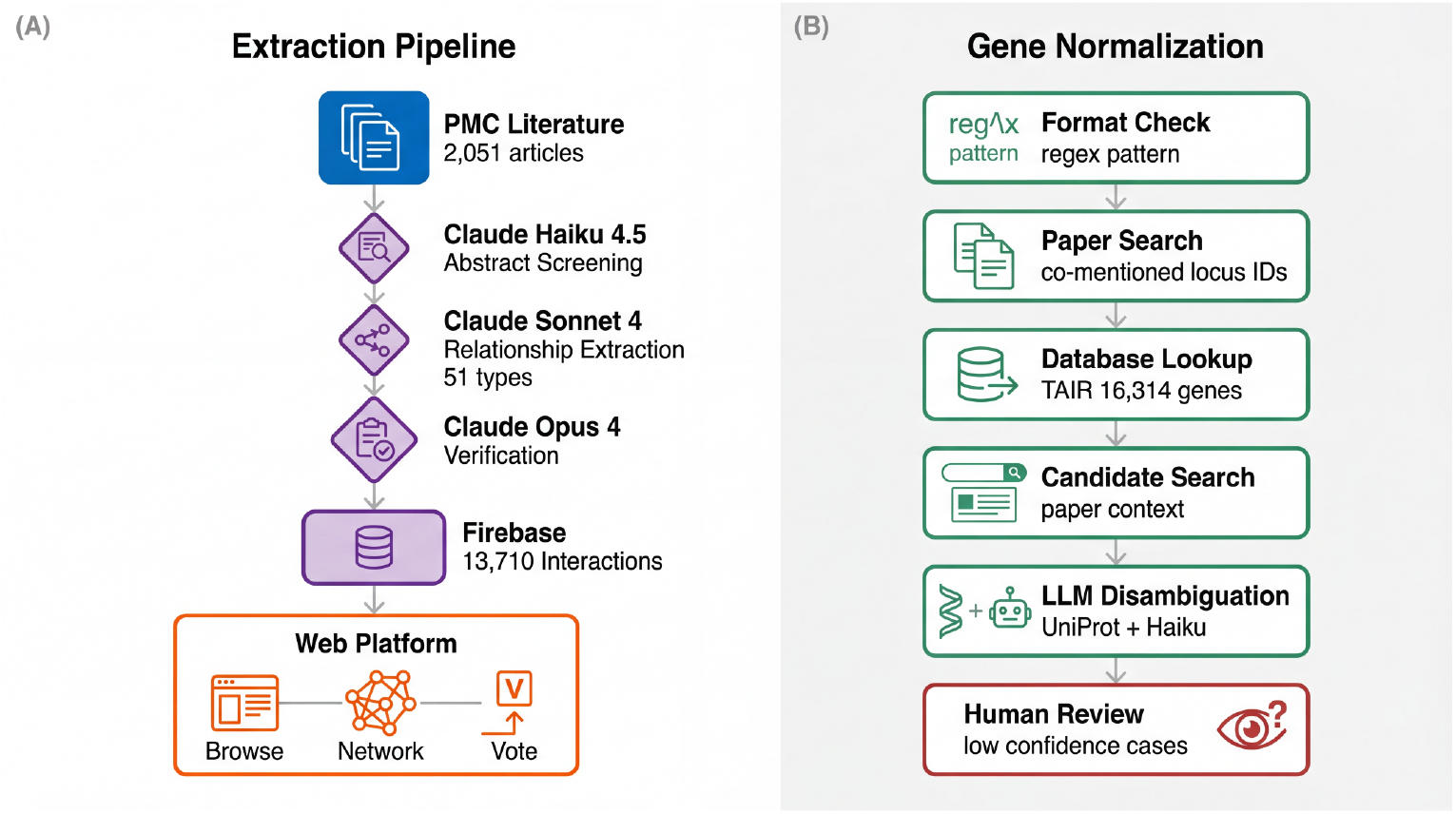
GeneReL Pipeline Architecture and Gene Normalization Workflow. (A) Overview of the data extraction and curation pipeline showing data collection from PMC, tiered AI processing (Haiku screening, Sonnet extraction, Opus verification), five-step gene normalization, and web application interfaces. (B) Five-step hierarchical gene normalization: format check, paper search for co-mentioned locus IDs, database lookup with alias support, paper search for candidates when ambiguous, and LLM disambiguation using UniProt annotations. Low-confidence cases flagged for human review.

- **Step 1 (Format Check):** Verify if the name already conforms to AGI locus ID format (AT#G#####).
- **Step 2 (Paper Search for Co-mentioned Locus IDs):** Search paper text for locus IDs co-mentioned near gene name occurrences within 50-character windows. This exploits the common practice of authors explicitly including identifiers (e.g., “FLC (AT5G10140)”).
- **Step 3 (Database Lookup):** Query the gene mapping dictionary containing 16,314 genes with alias support (**Supplementary Table S3**) [23], aggregated from TAIR and curator-validated mappings.
- **Step 4 (Paper Search for Candidates):** When multiple candidate locus IDs exist, search the paper for mentions of any candidate to resolve ambiguity using paper-specific context.
- **Step 5 (LLM Disambiguation):** For remaining ambiguous cases, invoke Claude Haiku 4.5 (claude-haiku-4-5-20251001) with UniProt functional annotations [24]. The model evaluates which candidate gene best matches the biological context described in the evidence text.

This paper-first strategy addresses the challenge that TAIR curators face when authors omit AGI identifiers, requiring “detective work to infer which identifier should be associated with the new gene symbol” [23]. Interactions where disambiguation yields Low confidence are automatically flagged for human review.

### 2.5 Web Platform Implementation

The web application was developed using React 18.2 with TypeScript and Tailwind CSS. Pages include: Home (card-based interaction browser with filtering), Network (Cytoscape.js visualization), My Votes (user voting history), and administrative Review/Suggestions interfaces (for only manager account).

Network visualization uses normalized locus IDs as unique node identifiers, ensuring genes mentioned under different names across papers consolidate into single nodes with correct topology (**Figure 2**). Node types include protein-coding genes (circles) and microRNAs (pink diamonds). Edge colors indicate relationship types, while thickness encodes confidence levels where multiple evidences increase the thickness.

**Figure 2.**
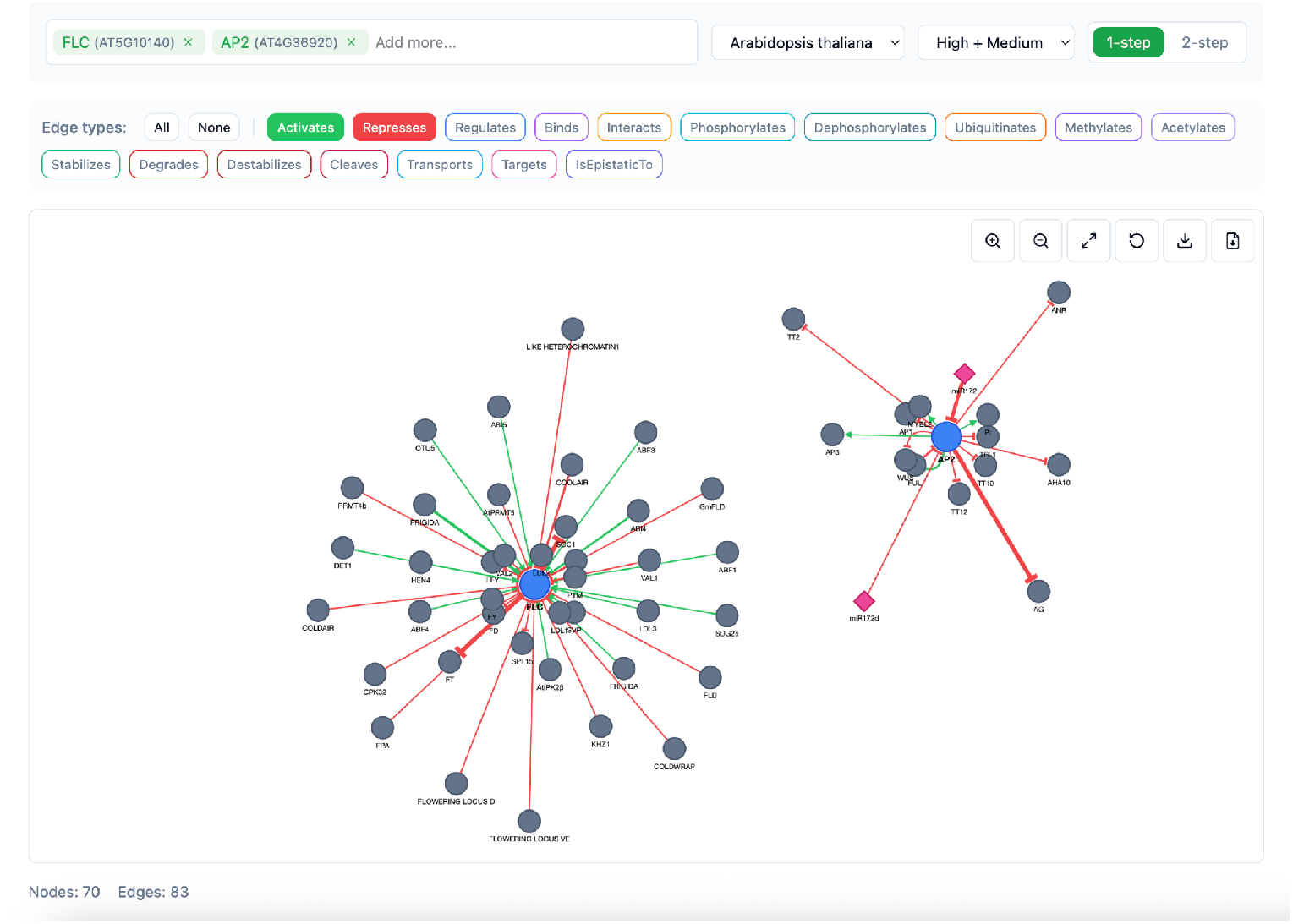
Network Visualization Features. Node types include protein-coding genes (circles) and microRNAs (pink diamonds). Edge colors indicate relationship types; thickness encodes confidence. Nodes use locus IDs as unique identifiers ensuring correct network topology.

### 2.6 Quality Control

Community curation through voting leverages collective expertise to identify high-quality versus questionable extractions [25]. Firebase Authentication manages user accounts via Google OAuth 2.0, with vote tracking employing atomic transactions for consistency.

## 3 Results

### 3.1 Database Composition

The GeneReL database contains 13,710 curated gene-gene interactions from *Arabidopsis thaliana*, classified into 51 distinct relationship types (**Supplementary Table S2**) supported by 9 evidence categories. Source publications span 2010-2024, with the current pipeline having processed 2,051 full-text articles from PubMed Central (**Supplementary Figure S2**). Publication years show peak coverage in 2018-2022, accounting for 54% of all extractions, reflecting both literature availability and the PMC open access corpus composition. Transcriptional regulation represents the predominant category (58.6%), dominated by Regulates (including Modulates and Controls; 24.4%), Activates (including Induces, Upregulates, Promotes, Enhances, and Enables; 18.7%), and Represses (including Suppresses, Inhibits, Downregulates, Reduces, Negatively Regulates, Restricts, and Blocks; 15.5%). Physical interactions (Interacts, Binds, and related types) account for 29.6%, while post-translational modifications (phosphorylation, ubiquitination, degradation) comprise 8.0%. Post-transcriptional regulation accounts for 2.8% of extracted interactions. (**Figure 3A and Supplementary Table S2**). The resulting interaction network is visualized on the web platform’s Network page using Cytoscape.js (**Figure 2**). Users can search for one or more genes and toggle between 1-step and 2-step network depth. Edge types can be filtered by relationship category. Clicking nodes displays all interactions for that gene, while clicking edges reveals supporting evidence sentences with links to source publications (**Supplementary Figure S4**).

**Figure 3.**
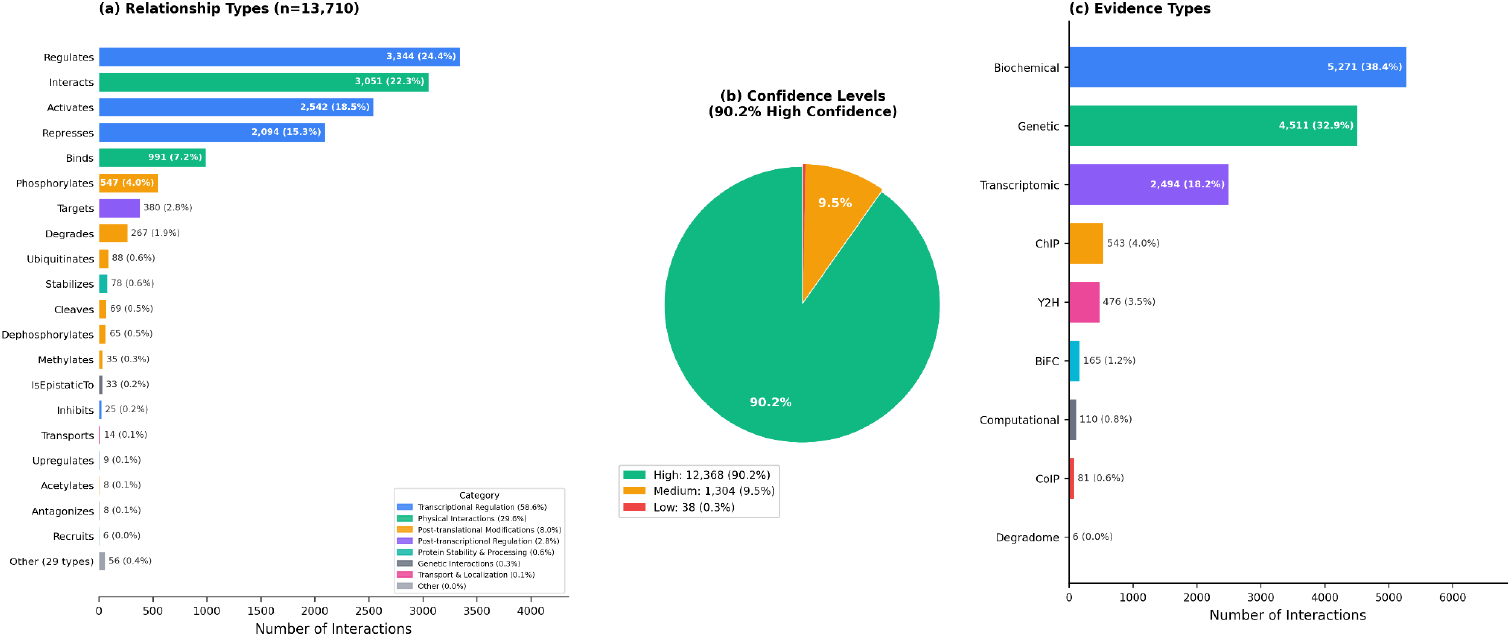
Database Composition. (A) Distribution of relationship types. (B) Confidence level distribution showing 90.2% high confidence. (C) Evidence type classification.

### 3.2 Confidence and Evidence Distribution

90.2% (12,368) of extracted relationships (13,710) received High confidence ratings, indicating definitive experimental demonstration (**Figure 3B**). Medium confidence interactions comprise 9.5% (1,304), reflecting qualified statements. Low confidence relationships account for 0.3% (38). Evidence type distribution: Biochemical (38.4%), Genetic (32.9%), Transcriptomic (18.2%), ChIP (4.0%), Y2H (3.5%), BiFC (1.2%), Computational (0.8%), CoIP (0.6%), and Degradome (<0.1%) (**Figure 3C**).

### 3.3 Gene Normalization Performance

The five-step pipeline achieves 50.7% successful gene normalization counted per interaction (**Supplementary Figure S5**). Step-by-step performance: Step 1 (Format Check) identified 3/13,710 (0.0%) interactions where genes were already in AGI locus format. Steps 2–4 (Dictionary & Paper Search, combining paper search for co-mentioned locus IDs, database lookup with alias support, and paper search for candidate locus IDs when ambiguous) resolved 6,260 (45.7%) of total interactions. miRNA Recognition resolved an additional 238 (1.7%). Step 5 (LLM-Assisted Resolution using UniProt functional annotations) resolved 445 (3.2%). Overall, 6,946/13,710 interactions (50.7%) were successfully normalized; 6,764 (49.3%) were flagged for human review or remained unresolved. Analysis of human review cases (n=7,223 interactions flagged with “needsReview”; unresolved + resolved but need to review) shows that normalization failure (not in common name to AGI locus format dictionary) accounts for 92.0%, LLM suggestions needing verification for 6.8%, ambiguous genes with multiple candidates for 0.7%, and other review needed for 0.5%.

### 3.4 Comparison with existing DB; STRING and IntAct

Of 13,710 GeneReL interactions, 6,569 (47.9%) have both genes resolved to AGI locus IDs and are thus matchable against external databases; the remaining interactions have one (28.9%) or neither (23.2%) gene resolved and cannot be meaningfully compared (**Figure 4a**). Among these 6,569 matchable interactions, 4,656 (70.9%) have corresponding gene pairs in STRING v12.5 high-confidence (score ≥ 700) interactions, while 1,913 (29.1%) are unique to GeneReL (**Figure 4b** and **Supplementary Table S4**) [18]. This high overlap validates the accuracy of GeneReL’s literature-derived extraction against an established interaction database. However, GeneReL’s unique contribution lies in its semantic richness: among the 1,913 interactions unique to GeneReL, regulatory relationships with directionality (Regulates: 516; Activates: 363; Represses: 296) go beyond STRING’s association-based scoring. Post-translational modifications (Phosphorylates: 39; Degrades: 20) and miRNA targeting (Targets: 42) capture regulatory mechanisms absent from STRING.

**Figure 4.**
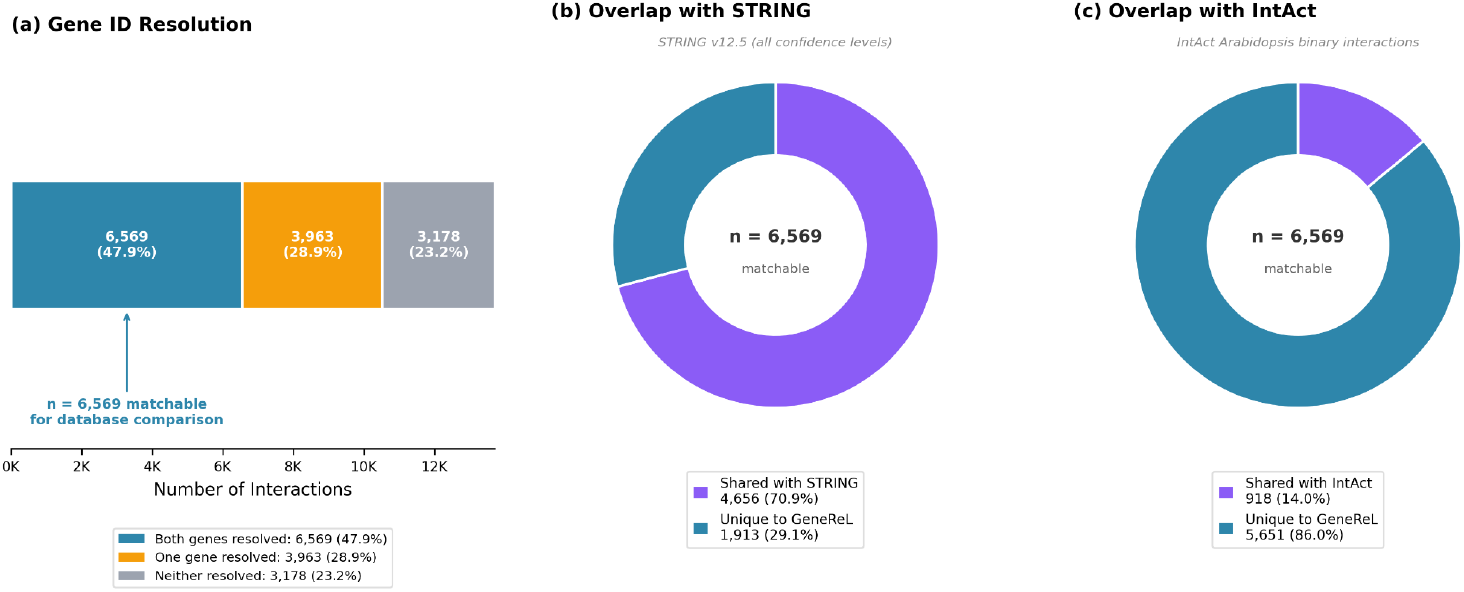
Comparison of GeneReL interactions with STRING and IntAct databases, restricted to matchable interactions where both genes are resolved to AGI locus IDs. **(a)** Gene ID resolution overview. Of 13,710 total GeneReL interactions, 6,569 (47.9%) have both genes resolved to AGI locus IDs and are matchable against external databases. The remaining interactions have one gene (28.9%) or neither gene (23.2%) resolved, and cannot be meaningfully compared. **(b)** Overlap of matchable GeneReL interactions (n = 6,569) with STRING v12.5 high-confidence (combined score >= 700) *Arabidopsis thaliana* interactions, determined by bidirectional gene-pair matching. 70.9% of matchable interactions are shared with STRING. **(c)** Overlap of matchable GeneReL interactions (n = 6,569) with IntAct Arabidopsis thaliana binary protein interactions. 14.0% of matchable interactions are shared with IntAct, reflecting IntAct’s focus on experimentally validated physical interactions versus GeneReL’s broader regulatory relationships from literature.

To further validate the unique contribution of literature-based extraction, we compared GeneReL interactions against the IntAct molecular interaction database [19], a primary curated resource for experimentally validated protein interactions. Gene pairs were matched bidirectionally (A-B equivalent to B-A; **Supplementary Table S4**). Of 6,569 matchable interactions, 5,651 (86.0%) represent unique relationships not present in IntAct, while 919 (14.0%) have corresponding gene pairs (**Figure 4c**). The high proportion of unique interactions reflects fundamental differences between databases: IntAct primarily captures physical associations from high-throughput screens (Y2H, mass spectrometry, co-immunoprecipitation), while GeneReL extracts semantically rich regulatory relationships from targeted experimental studies described in narrative text. Notably, relationship types with no IntAct equivalent include Regulates (1,352 unique), Interacts (1,180 unique), Activates (1,137 unique), Represses (958 unique), and miRNA Targets (118 unique).

### 3.5 Validation

Manual validation of 100 randomly selected high-confidence extractions from the GeneReL database was performed. Each extraction was evaluated on three criteria: (1) correct gene identification—whether both gene names appear in the supporting evidence sentence; (2) appropriate relationship type—whether the evidence sentence contains language consistent with the assigned interaction type; and (3) evidence relevance—whether the supporting sentence is a well-formed, meaningful text passage containing at least one gene name. Results confirm approximately 90% of extractions accurately reflect source literature claims (**Supplementary Table S7**). However, several cases of validation failure resulted from sentence parsing errors while the gene-to-gene relationship itself was correct (e.g., “APRF1 as a Repressor of FLC Expression” extracted correctly but with truncated evidence text). Some unpredictable parsing errors produce incomplete sentences in the evidence field; however, as the pipeline continuously adds evidence from new publications, multiple supporting sentences accumulate for each interaction edge. This redundancy allows users to cross-validate relationships and identify problematic extractions through the voting mechanism, flagging incorrect interaction cards for future curator correction. Comparison with well-characterized pathways confirms capture of canonical relationships including FLC repression of FT, CO activation of FT, and SVP repression of SOC1.

Network hub analysis reveals coverage biases reflecting publication patterns in the source literature. Well-studied regulatory genes (e.g., FLC, CO, AG, AP1, ABI3) are substantially over-represented: the top 50 most-connected nodes account for 24% of total network degree (**Supplementary Table S6**). This reflects both genuine biological importance of these regulatory hubs and systematic under-representation of less-studied genes.

## 4 Discussion

### 4.1 Principal Findings

This work demonstrates successful integration of LLM-based extraction with community curation for biological database construction. The LLM extraction achieves high precision in recognizing diverse relationship types across multiple regulatory layers.

Ablation analysis confirms the value of multi-step normalization: format check alone (Step 1) resolved only a bit (∼0.0%) of interactions, while Dictionary & Paper Search (Steps 2–4) contributed the largest gain at 45.7%. miRNA Recognition added 1.7%, and LLM-Assisted Resolution (Step 5) contributed an additional 3.2%, bringing the full pipeline to 50.7% successful normalization. The remaining 49.3% of interactions were flagged for human review, reflecting the inherent complexity of gene nomenclature disambiguation. This performance is consistent with advances in gene normalization systems such as GNorm2 [26], though direct comparison is complicated by different evaluation corpora.

Complementary coverage with existing databases is demonstrated by comparison with IntAct, where 86.0% of matchable GeneReL interactions represent unique gene pairs not present in that curated resource. This substantial difference reflects the fundamentally distinct data sources: IntAct aggregates high-throughput experimental data capturing physical associations, while GeneReL extracts semantically rich regulatory relationships from narrative descriptions of targeted experiments. The complementarity validates GeneReL’s contribution to the interaction database ecosystem rather than redundancy with existing resources.

### 4.2 LLM Advantages and Limitations

LLMs demonstrate advantages over earlier text mining methodologies. The extensive pre-training provides robust handling of nomenclature heterogeneity, while contextual understanding enables extraction across linguistic variation without exhaustive rule specification. Zero-shot to few-shot learning proves valuable for specialized domains with limited annotated corpora. However, LLMs occasionally generate plausible but incorrect extractions—hallucinations that cannot occur with rule-based systems. The human-in-the-loop correction strategy addresses this by routing Low-confidence disambiguations to curator review.

### 4.3 Community Curation

The voting mechanism implements scalable quality assessment, distributing validation workload across the research community following successful precedents [5,25]. Maintaining transparent display of both positive and negative vote counts provides users with agreement information while preserving access to all extracted data. Early adopter feedback indicates particular utility for flowering time and hormone signaling pathway researchers.

### 4.4 Limitations

The extraction operates on individual passages independently, lacking mechanisms to integrate evidence across papers or resolve conflicting claims. Gene normalization remains imperfect for recently characterized genes lacking UniProt entries. The current schema captures relationship existence but loses contextual detail including tissue-specificity and developmental stage dependence. Coverage biases inherent to literature-derived databases mean users should interpret network centrality metrics with awareness of publication patterns rather than assuming uniform genome-wide coverage.

## 5 Conclusion

GeneReL demonstrates successful integration of modern language models with community curation for biological database construction. By combining LLM extraction providing scalable processing with distributed validation ensuring quality control, the system addresses longstanding challenges in knowledge resource development. The platform provides the *Arabidopsis* research community with an accessible, continuously updated repository of regulatory relationships with transparent evidence provenance, advancing efforts to transform biomedical literature into structured knowledge resources.

## Author Contributions

**Joo-Seok Park:** Conceptualization, Methodology, Software, Writing - Original Draft, Visualization. **Su-Yeon Ha:** Data Curation, Validation. **Yejin Lee:** Software (Cloud Infrastructure). **Yang Jae Kang:** Supervision, Project Administration, Funding Acquisition, Writing - Review & Editing.

## Data Availability

The GeneReL database and web platform are freely accessible at https://generel.newgenes.me. Data can be downloaded in CSV and JSON formats.

