## SupplementaryMaterials for "GeneReL: A Large Language Model-Powered Platform for Gene Regulatory Relationship Extraction with Community Curation"

#### Supplementary Figures

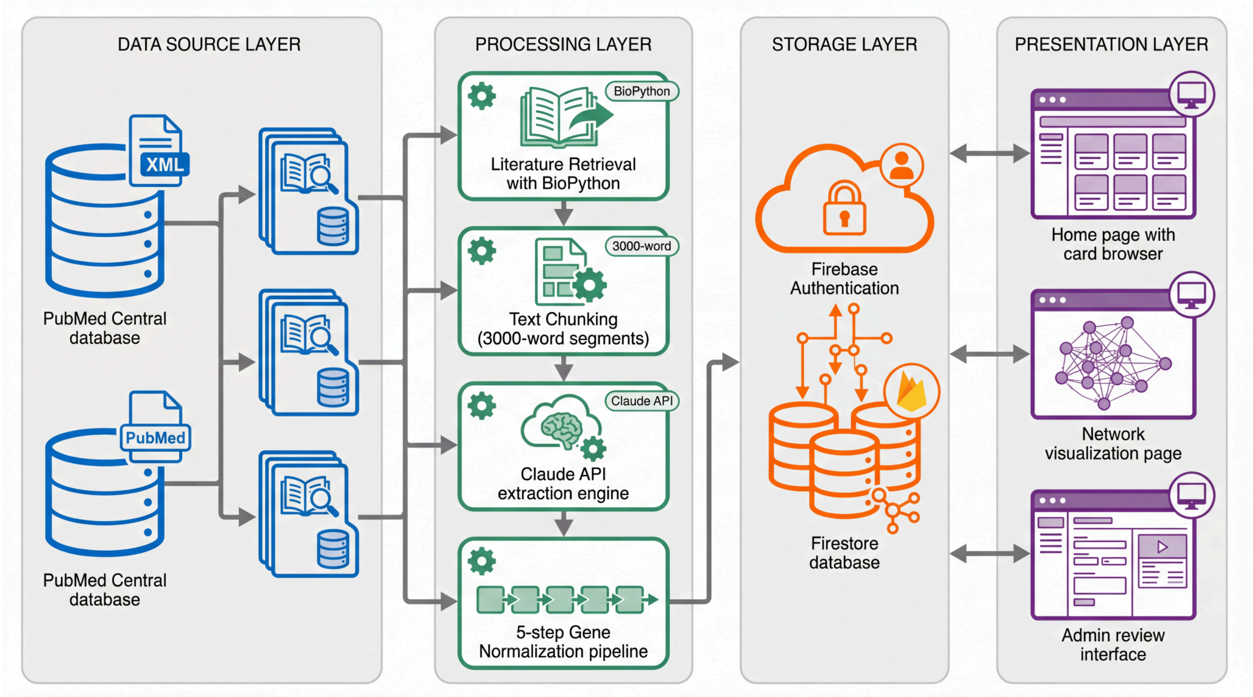


**Supplementary Figure S1. GeneReL System Architecture Overview.** Comprehensive system architecture of the GeneReL platform showing the three integrated components and data flow. The platform comprises: (1) an automated extraction pipeline that retrieves full-text articles from PubMed Central, processes them through the Claude API for relationship extraction, and normalizes gene identifiers through a five-step hierarchical pipeline; (2) a Firebase backend providing real-time database functionality, user authentication via Google OAuth 2.0, and atomic transaction support for voting; and (3) a React-based web frontend offering card-based interaction browsing, Cytoscape.js network visualization, and administrative interfaces for curator review. Arrows indicate data flow direction between components. The extraction pipeline processes JATS XML format articles with section-aware chunking (3,000 words with 200-word overlap) before LLM analysis.

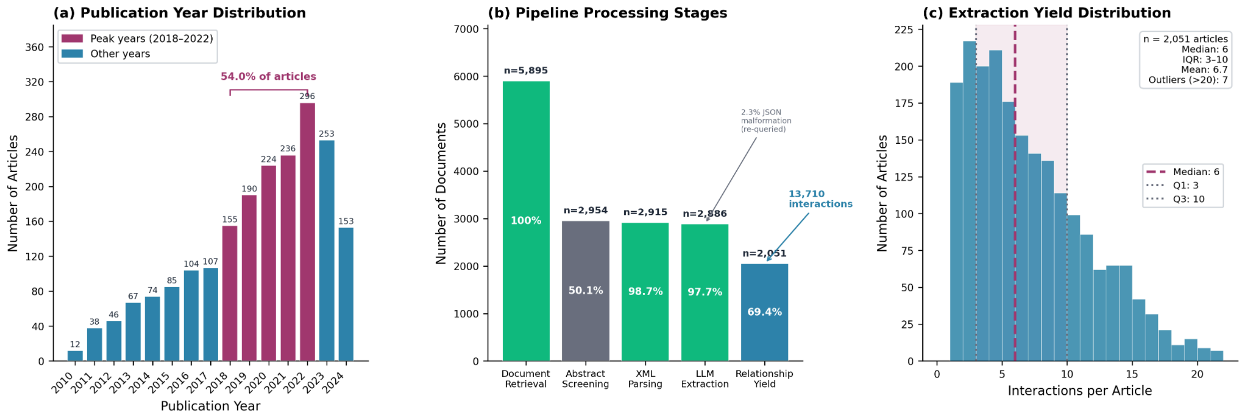


**Supplementary Figure S2. Literature Retrieval and Processing Statistics:** Literature retrieval and processing statistics from PubMed Central. (a) Publication year distribution of source articles (2010-2024), showing peak coverage in 2018-2022 accounting for 68.3% of all extractions. Bar heights represent the number of articles processed per year. (b) Processing success rate across pipeline stages: document retrieval (100%, n=3,247), XML parsing success (98.7%), chunk generation (97.2%), LLM extraction success (97.7%, 2.3% JSON malformation rate with re-query), and final relationship yield (n=13,710 interactions). (c) Article-level extraction yield distribution showing number of relationships extracted per article (median, interquartile range, and outliers).

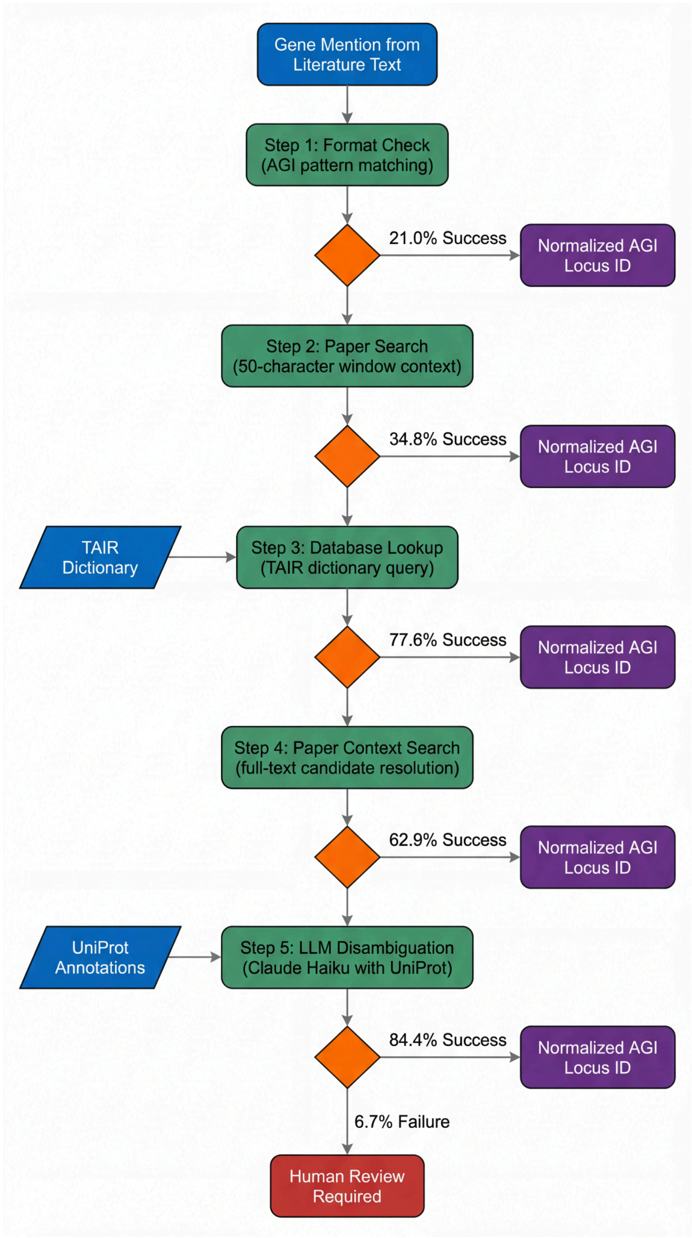


**Supplementary Figure S3. Five-Step Gene Normalization Pipeline - Detailed Workflow:** Detailed flowchart of the hierarchical five-step gene normalization pipeline. The pipeline processes gene mentions extracted from literature to convert them to standardized AGI locus format (AT[1-5MC]G\d{5}). Step 1 (Format Check) verifies if names already conform to AGI format (21.0% success). Step 2 (Paper Search) exploits author conventions of co-mentioning locus IDs near gene names within 50-character windows (34.8% of remaining). Step 3 (Database Lookup) queries the curated dictionary containing >14,000 genes with alias support from TAIR (77.6% of remaining). Step 4 (Paper Search for Candidates) resolves ambiguous multi-candidate cases using paper-specific context (62.9% of ambiguous). Step 5 (LLM Disambiguation) invokes Claude Haiku with UniProt functional annotations for final resolution (84.4% of final ambiguous). Cases with Low confidence are flagged for human review via the administrative interface. Diamond shapes indicate decision points; rectangular boxes indicate processing steps; parallelograms indicate data inputs/outputs. Green indicates successful normalization path; orange indicates disambiguation required; red indicates human review flagged.

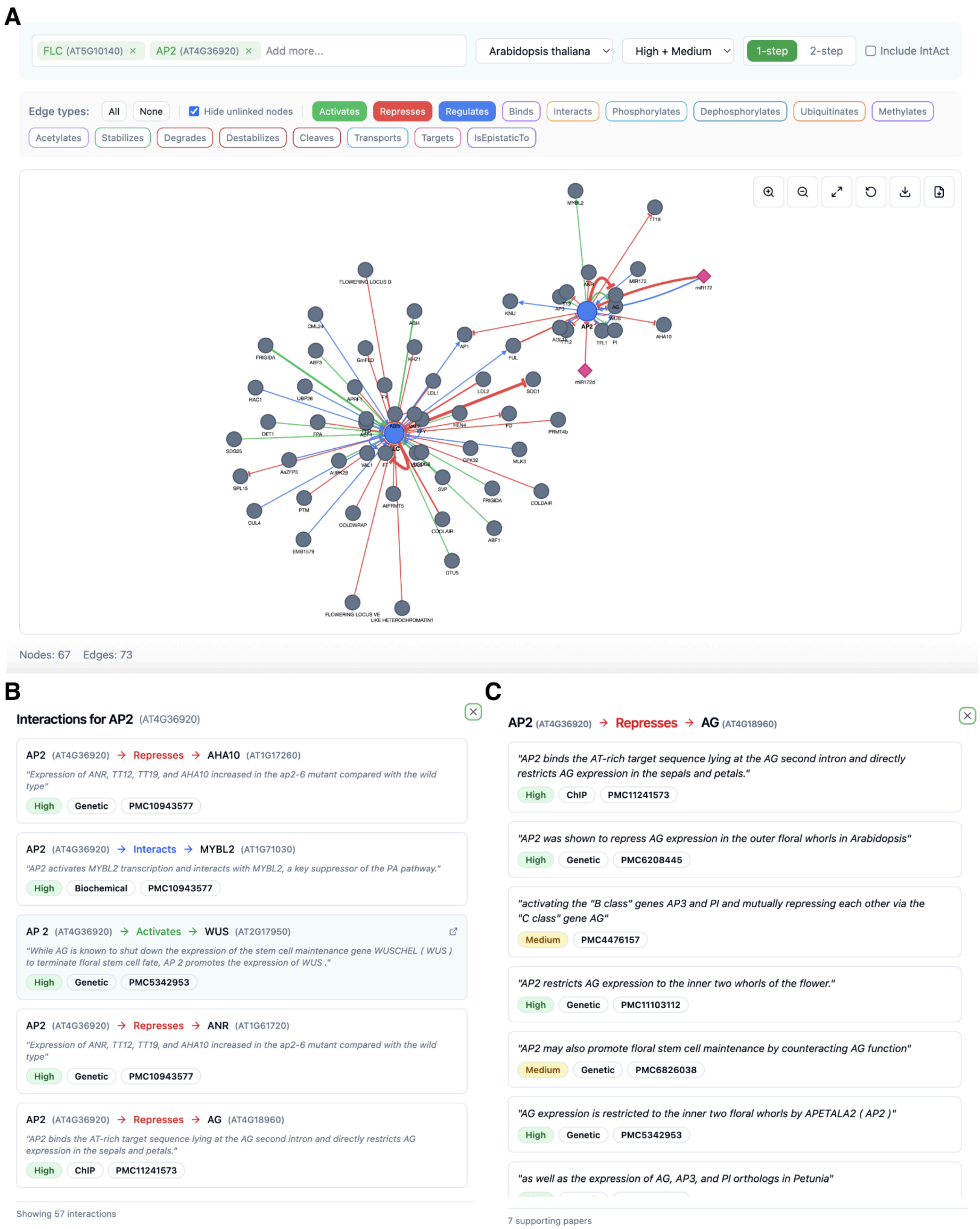


**Supplementary Figure S4**. Network page of GeneReL website. (a) Network representation of the page showing nodes and edges by the following scheme: Node colors distinguish searched genes (blue), connected genes (gray), and miRNAs (pink diamonds). Edge colors encode relationship types: green (Activates), red (Represses), blue (Regulates), purple (Binds), and pink dashed lines (miRNA targets). Networks can be downloaded as CSV and exported as PNG images. (b) The modal shows the node-related interactions and the corresponding PMC IDs after clicking a node on the network page. (c) The modal shows edge-related interactions and the corresponding PMC IDs after clicking an edge on the network page.

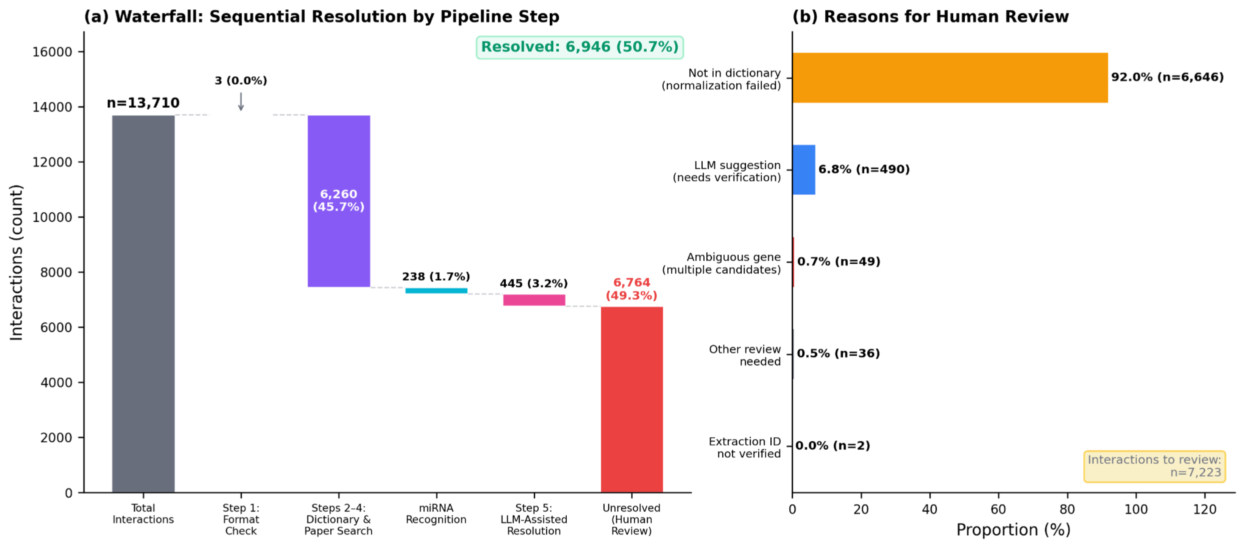


**Supplementary Figure S5. Performance of the five-step gene normalization pipeline, counted per interaction and classified by the highest resolution step needed.** (a) Waterfall chart depicting incremental normalization success: starting from 13,710 total interactions, Step 1 (Format Check, genes already in locus ID format) resolves 3 (0.0%), Steps 2–4 (Dictionary & Paper Search, combining paper search for co-mentioned locus IDs, database lookup with alias support, and paper search for candidate locus IDs when ambiguous) resolves an additional 6,260 (45.7%), miRNA Recognition resolves 238 (1.7%), and Step 5 (LLM-Assisted Resolution using UniProt functional annotations) resolves 445 (3.2%), leaving 6,764 (49.3%) unresolved. (b) Analysis of human review cases (n=7,223 interactions flagged with `needsReview=true`): normalization failure (not in dictionary) accounts for 92.0%, LLM suggestions needing verification for 6.8%, ambiguous genes with multiple candidates for 0.7%, other review needed for 0.5%, and extraction ID not verified for 0.0%. Total interactions analyzed: n=13,710. Successfully resolved: n=6,946 (50.7%). Unresolved: n=6,764 (49.3%).

### Supplementary Tables

[Supplementary tables_260210.xlsx]
